# LRRK2/LRRK1 interactions modulate Rab7 activity and inhibit lysosomal exocytosis

**DOI:** 10.64898/2026.06.12.731951

**Authors:** Madiha Merghani, Ellen Gerhardt, Maiken Hesse, Christiane Fahlbusch, C. Alexander Böcker, Tiago F. Outeiro

**Affiliations:** University Medical Center Göttingen, Department of Experimental Neurodegeneration, Center for Biostructural Imaging of Neurodegeneration, Germany; University Medical Center Göttingen, Department of Neurology, Germany; Faculdade de Medicina e Ciências Biomédicas, Algarve Biomedical Center Research Institute (ABC-Ri), Universidade do Algarve, 8005-139 Faro, Portugal; Translational and Clinical Research Institute, Faculty of Medical Sciences, Newcastle University, UK; Deutsches Zentrum für Neurodegenerative Erkrankungen (DZNE), Göttingen, Germany

**Keywords:** Parkinsońs disease, LRRK2, LRRK1, Lysosome, autophagy

## Abstract

Mutations in the *LRRK2* gene are the most common genetic cause of both familial and sporadic Parkinson’s disease (PD). LRRK2 belongs to the leucine-rich repeat kinase (LRRK) family. Two members of the LRRK family exist in humans (LRRK1 and LRRK2). Although there is strong structural similarity between the two proteins, they have attracted very different levels of attention by the scientific community owing to the strong association between LRRK2 and PD. In contrast, the role of LRRK1 is relatively unexplored. LRRK2 is also known to regulate endolysosomal function, but its precise role in this process remains incompletely understood. Our study investigated the interaction between LRRK1 and LRRK2 under different cellular conditions, uncovering their role in modulating the endolysosomal system. We found that LRRK1 and LRRK2 interact and modulate each other’s activity, and that this interaction is reduced under starvation conditions. We also found that LRRK1 and LRRK2 have contrasting effects on lysosomal size, impacting on lysosomal exocytosis. Together, our findings suggest that LRRK2 regulates endolysosomal homeostasis, at least in part, by modulating LRRK1. Our findings offer new insight into the molecular mechanisms associated with lysosomal function and, ultimately, we anticipate this knowledge will help us better understand the molecular crosstalk between LRRK kinases and their contribution to PD pathogenesis.

**Graphical abstract:** Starvation reduces the interaction between LRRK2 and LRRK1 due to a conformational change in LRRK2. Under normal conditions, LRRK2/LRRK1 interaction enhances LRRK1 activity, leading to increased phosphorylation of Rab7. Disruption of the Rab7 cycle impairs lysosomal homeostasis, leading to lysosomal accumulation and an increase in lysosomal diameter. This enlargement negatively impacts lysosomal exocytosis. Created with BioRender.com.

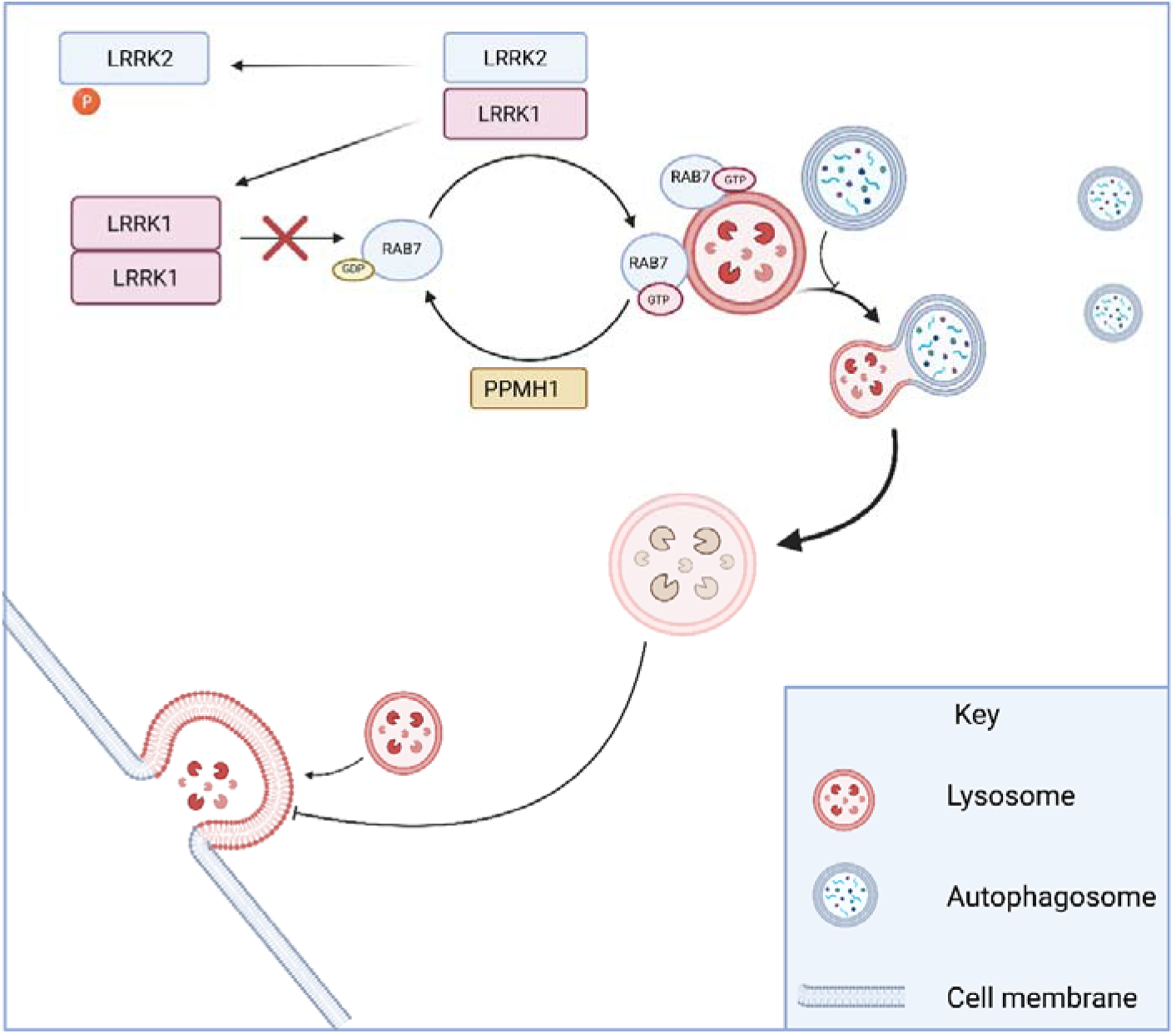

## Introduction

PD is a complex and progressive neurodegenerative disorder in which multiple factors converge to drive the selective degeneration of substantia nigra dopaminergic neurons. In addition to neuronal loss, the accumulation of pathognomonic intraneuronal α-synuclein (aSyn) inclusions known as Lewy bodies is a key hallmark of the disease (1–3). However, the mechanisms underlying aSyn accumulation and dopaminergic neuron degeneration remain unclear, limiting our ability to rationally develop effective disease-modifying treatments (DMTs).

The autophagy–lysosomal pathway (ALP) is one of the most studied pathways in PD, not only because it is the primary pathway for aSyn degradation but also because the lysosomal dysfunction is a prominent feature of the disease (4–6). Moreover, several PD-linked genes are associated with lysosomal function, including *GBA*, *LRRK2*, and *VPS35* (7). Consistently, multiple studies highlight the significant role of LRRK2 in regulating lysosomal biology. LRRK2 influences lysosomal distribution, pH, morphology, degradative capacity, and the expression of lysosomal proteins. LRRK2 is also recruited to and activated on stressed lysosomes (8–10), sensing lysosomal membrane damage, and plays an important role in lysosomal remodelling and repair (11–14). Upon lysosomal damage, LRRK2 phosphorylates Rab8, thereby promoting the activation of the ESCRT-III machinery via recruitment of CHMP4B to the surface of the membrane to facilitate membrane repair (11). Another essential lysosomal process potentially regulated by LRRK2 is lysosomal exocytosis (15,16). This process is essential for plasma membrane repair and remodelling, ATP and proton release, and immune responses, regulating antigen presentation (17,18). LRRK2 phosphorylates Rab8A, Rab10, and Rab3A, all of which are involved in membrane trafficking, including exocytosis (19).

Another key regulator of lysosomal activity is Rab7. Rab7 is not a substrate of LRRK2 but of LRRK1, a protein within the same LRRK family (19). LRRK1 shares structural similarities with LRRK2, and these similarities extend to their biological functions and tissue expression patterns. Both proteins regulate intracellular trafficking through phosphorylation of Rab GTPases, facilitating cargo transport along microtubules. LRRK proteins are expressed in the brain, lung, heart, and kidney, in addition to other tissues as well (20–23). Despite these similarities, there are clear distinctions between LRRK1 and LRRK2. First, they phosphorylate different subsets of Rab proteins. Second, mutations in these proteins are associated with distinct diseases: LRRK2 mutations are linked to PD, whereas LRRK1 mutations are associated with a rare bone disease. Finally, LRRK1 mutations are thought to result in loss of function, whereas PD-associated LRRK2 mutations are considered gain-of-function (19,23–27). Adding to that, homodimerization of both LRRK1 and LRRK2 is well established phenomena (28–30). However, their functional outcomes differ: LRRK2 homodimerization enhances kinase activity and promotes microtubule association, whereas LRRK1 homodimerization traps the kinase domain in an inactive conformation (29,31). With all the advances in our understanding of LRRK protein structure and function, it remains unclear how these proteins are regulated within the cell and whether they cooperate in controlling different cellular processes. Here, we investigated the interaction between these two proteins revelling the cellular condition affecting their interaction. In addition, we investigated how this interaction changes the activity of LRRK proteins, leading to changes in lysosomal exocytosis.

## Results

### LRRK2 interacts with LRRK1

First, we investigated whether LRRK1 can interact with LRRK2. To do this, we used a proximity ligation assay (PLA), and found a significant increase in PLA puncta in cells co-expressing wild-type LRRK1 (WT LRRK1) and wild-type LRRK2 (WT LRRK2), compared with cells that do not express LRRK2 (Fig. 1A and B). Notably, this interaction persisted in cells expressing the LRRK2 G2019S mutant or the kinase-dead (KD) variant (Fig. 1A and B), indicating that the interaction occurs independently of LRRK2 kinase activity. The PLA signal was broadly distributed throughout the cytoplasm, suggesting that the LRRK1/LRRK2 interaction takes place in the cytosol. To study the structural basis of this interaction, we used AlphaFold 2 to predict the 3D structures of LRRK1 and LRRK2 and assess potential interaction interfaces. The predicted structures revealed that LRRK1 and LRRK2 align in a parallel orientation, with their COR domains forming the interaction interface (Fig. 1C and D). This mode of interaction closely resembles the known parallel homodimeric arrangement of LRRK2 (29).

**Fig 1.**
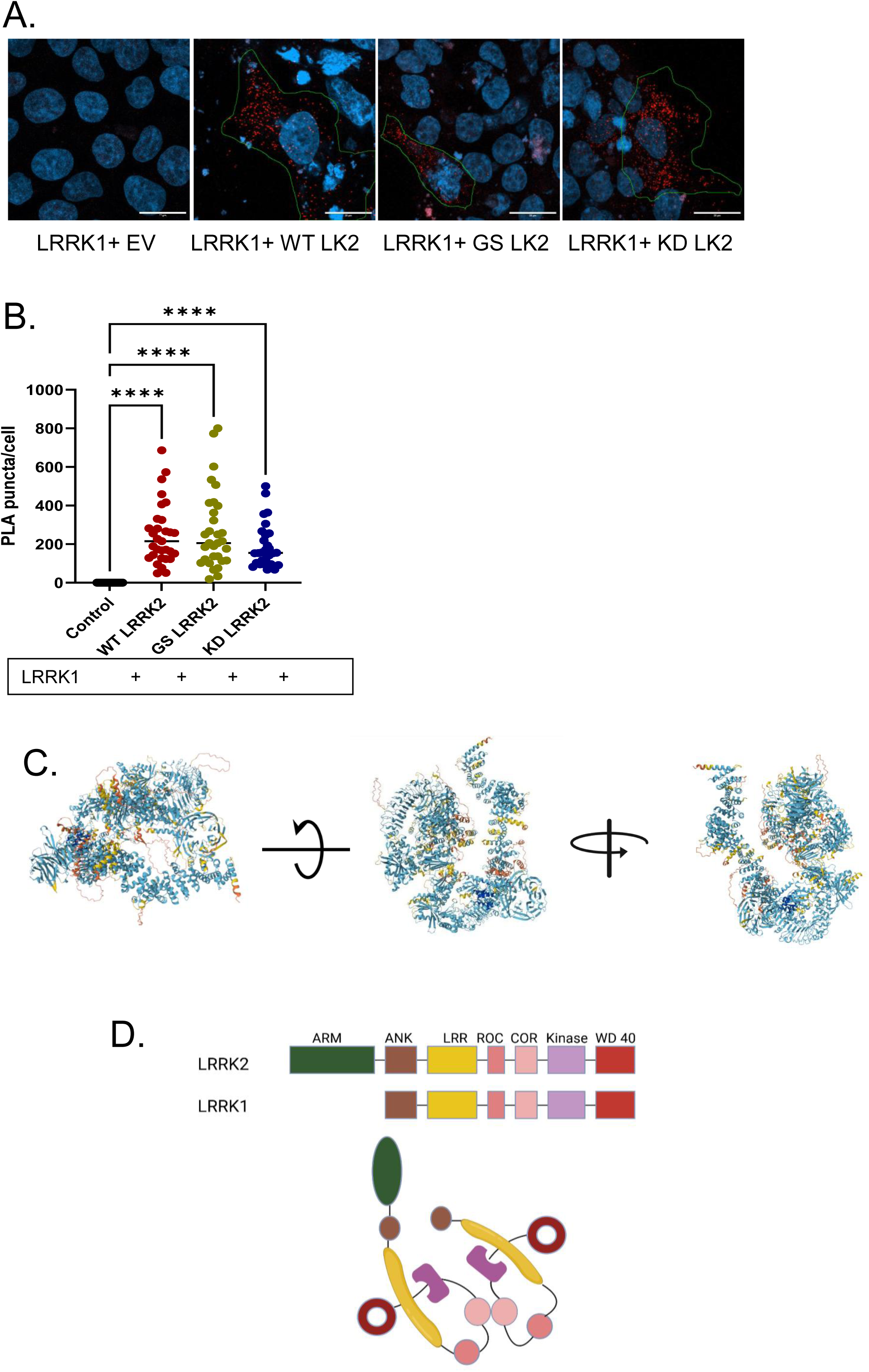
LRRK1 interacts with LRRK2. (**A**) Representative confocal images showing the PLA signal in red and the nucleus in blue in a cell transfected with LRRK1 and a different variant of LRRK2. (**B**) Quantification of the PLA signal (n=3; 1-way ANOVA; Dunnett’s multiple comparisons test, **** p<0.0001). (**C**) Alpha fold structure prediction of LRRK1/LRRK2 interaction. (**D**) Illustration of LRRK1/LRRK2 different domains and the possible interaction location based on alpha fold structure.

### LRRK2 / LRRK1 interaction is reduced under autophagic conditions in a kinase-dependent manner

To investigate the interaction between LRRK1 and LRRK2 in living cells, we employed the biomolecular fluorescence complementation (BiFC) system. We co-transfected cells with LRRK1 and LRRK2 fused to complementary fragments of a fluorescent reporter protein (Venus fluorescent protein). We observed a signal in cells co-expressing both LRRK1 and LRRK2, confirming their interaction (Fig. 2A). Notably, while the LRRK2 homodimer signal remained unchanged under starvation conditions, the LRRK2/LRRK1 BiFC signal significantly decreased (Fig. 2A-D). This suggests that starvation induces a conformational change that disrupts the LRRK2/LRRK1 interaction. The reduction in BiFC signal was more pronounced in cells expressing the LRRK2 G2019S mutant (Fig. 3A-D), indicating that the reduction in interaction between the two proteins is dependent on LRRK2 kinase activity. Interestingly, in these cells, the BiFC signal did not distribute diffusely in the cytoplasm but instead formed distinct, dot-like structures. The change in the BiFC signal with starvation led us to hypothesize that starvation alters the conformation of LRRK2, thereby modulating its interaction with LRRK1. Since phosphorylation at S935 is a well-established marker of LRRK2 conformational state, we assessed S935 phosphorylation under both optimal nutrient and starved conditions. We found that S935 phosphorylation increased significantly upon starvation, while total LRRK2 levels remained unchanged (S1. A, B, and D). This supports our hypothesis that starvation induces a conformational change in LRRK2, which disrupts its interaction with LRRK1.

**Fig 2.**
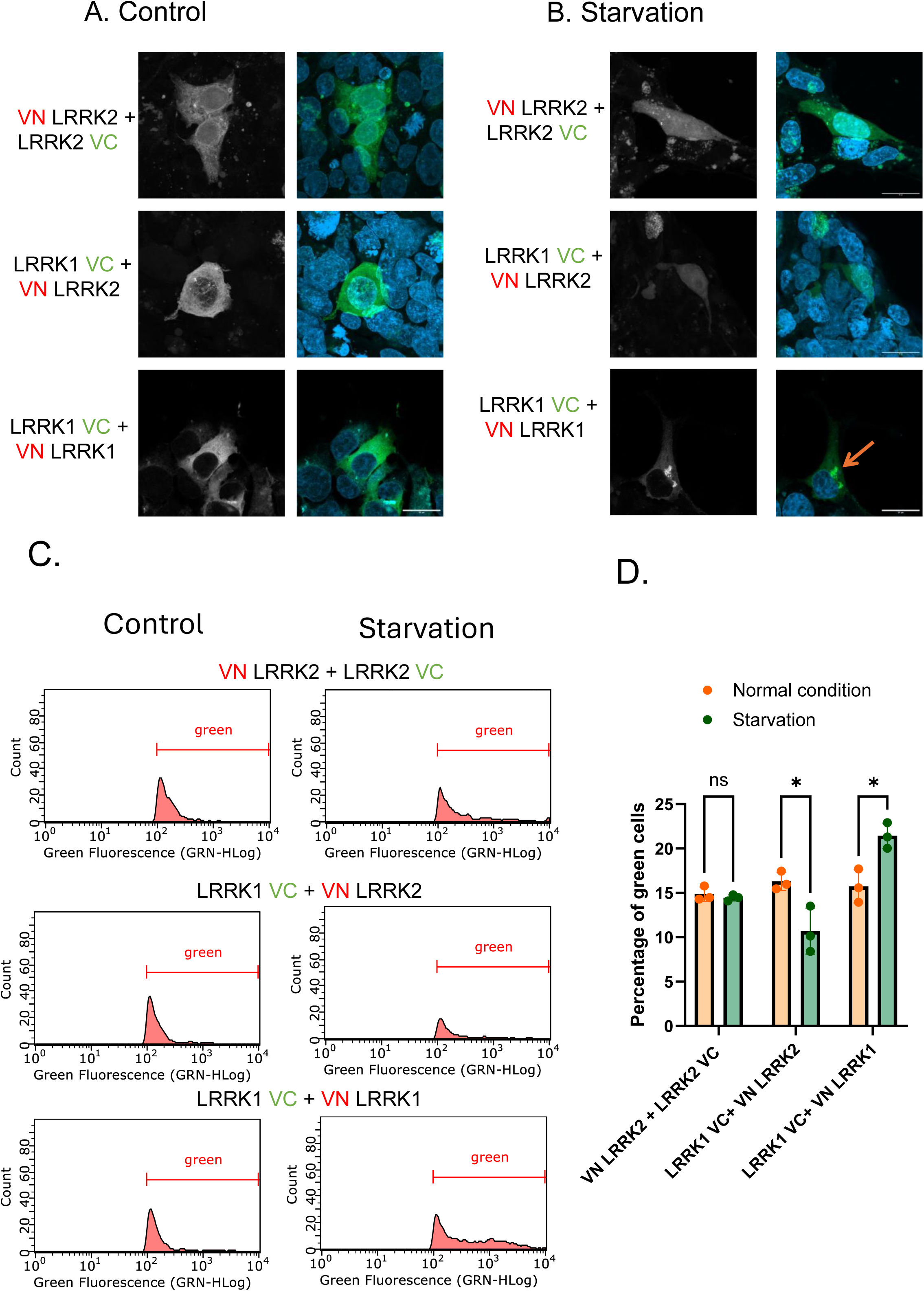
LRRK2/LRRK1 interaction decreases under autophagic conditions, while LRRK1 homodimer formation increases. (**A**) Representative confocal images showing the BiFC signal in HEK 293T under normal and (**B**) starvation conditions. (**C**) Flow cytometry histograms show the percentage of green cells under normal (left) and starvation conditions (right). (**D**) Date represents the percentage of green cells (n=3, Two-way ANOVA; Šídák’s multiple comparisons test, p= 0,0406 and 0,0387).

**Fig 3.**
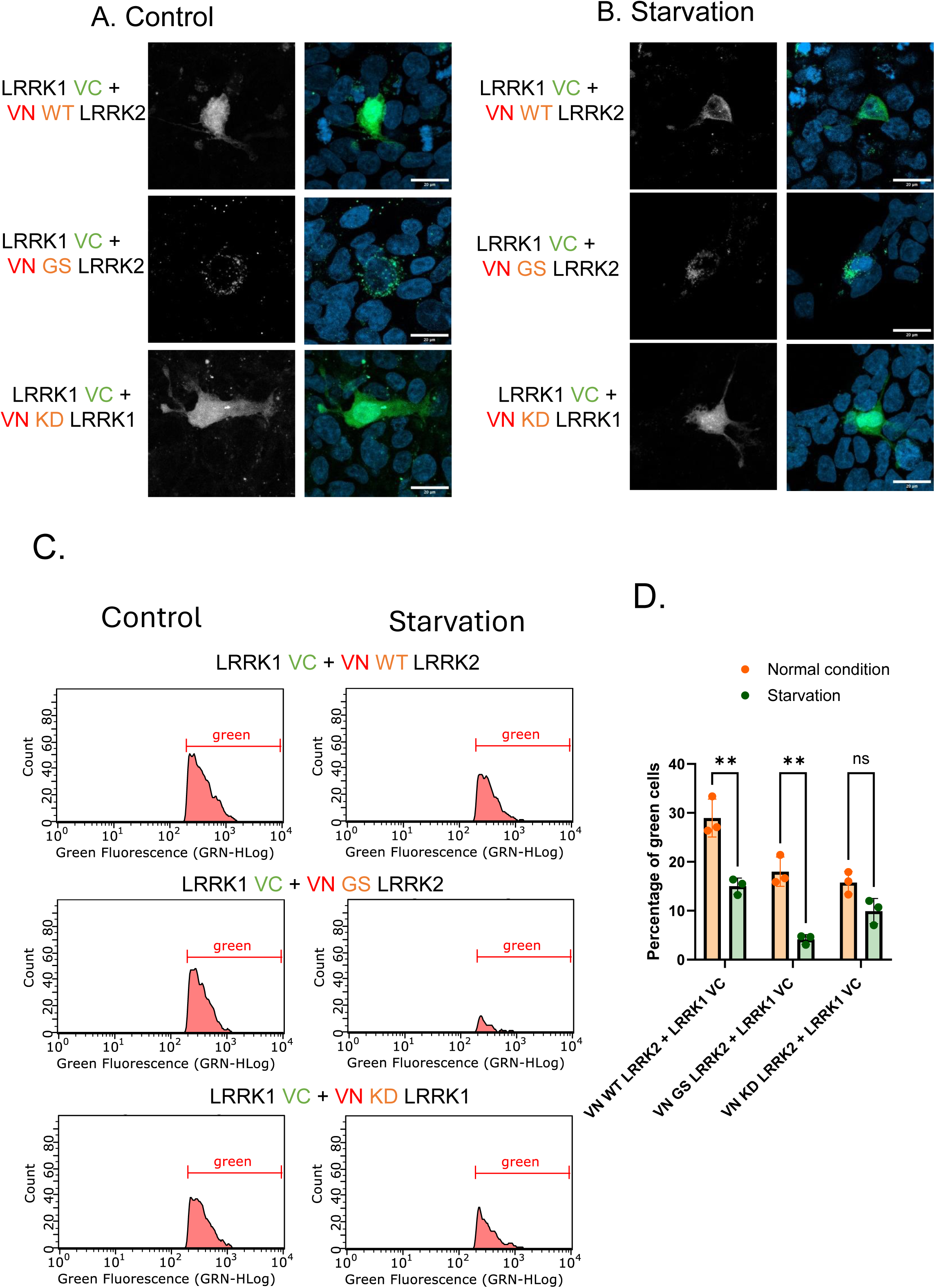
LRRK2/LRRK1 interaction decreases in a kinase-dependent manner under autophagic conditions. (**A**) Representative confocal images showing the BiFC signal in HEK 293T under normal and (**B**) starvation conditions. (**C**) Flow cytometry histograms show the percentage of green cells under normal (left) and starvation conditions (right). (**D**) Date represents the percentage of green cells (n=3, Two-way ANOVA; Šídák’s multiple comparisons test, p= 0,0012 and p=0,0012)

### phosphorylation of LRRK2 at S935 is reduced by LRRK1 under starvation conditions

Next, we investigated the functional impact of LRRK1 on LRRK2 kinase activity in cells co-expressing both proteins. Since phosphorylation at S935 is a well-established marker of LRRK2 conformation and is indirectly linked to its kinase activity, we used it as an indicator for LRRK2 activity. S935 phosphorylation is reduced upon treatment with type I LRRK2 kinase inhibitors, making it a possible indicator of kinase activity. To assess the effect of LRRK1 on LRRK2, we co-expressed WT LRRK2, G2019S, and KD mutant with WT LRRK1 in HEK293T cells. We found that LRRK1 significantly reduced S935 phosphorylation in WT LRRK2 and G2019S (Fig 4. A and B). This indicates that LRRK1 suppresses LRRK2 kinase activity. These results suggest that the LRRK1/LRRK2 interaction induces a conformational change and stabilizes LRRK2 inactive conformation.

**Fig 4.**
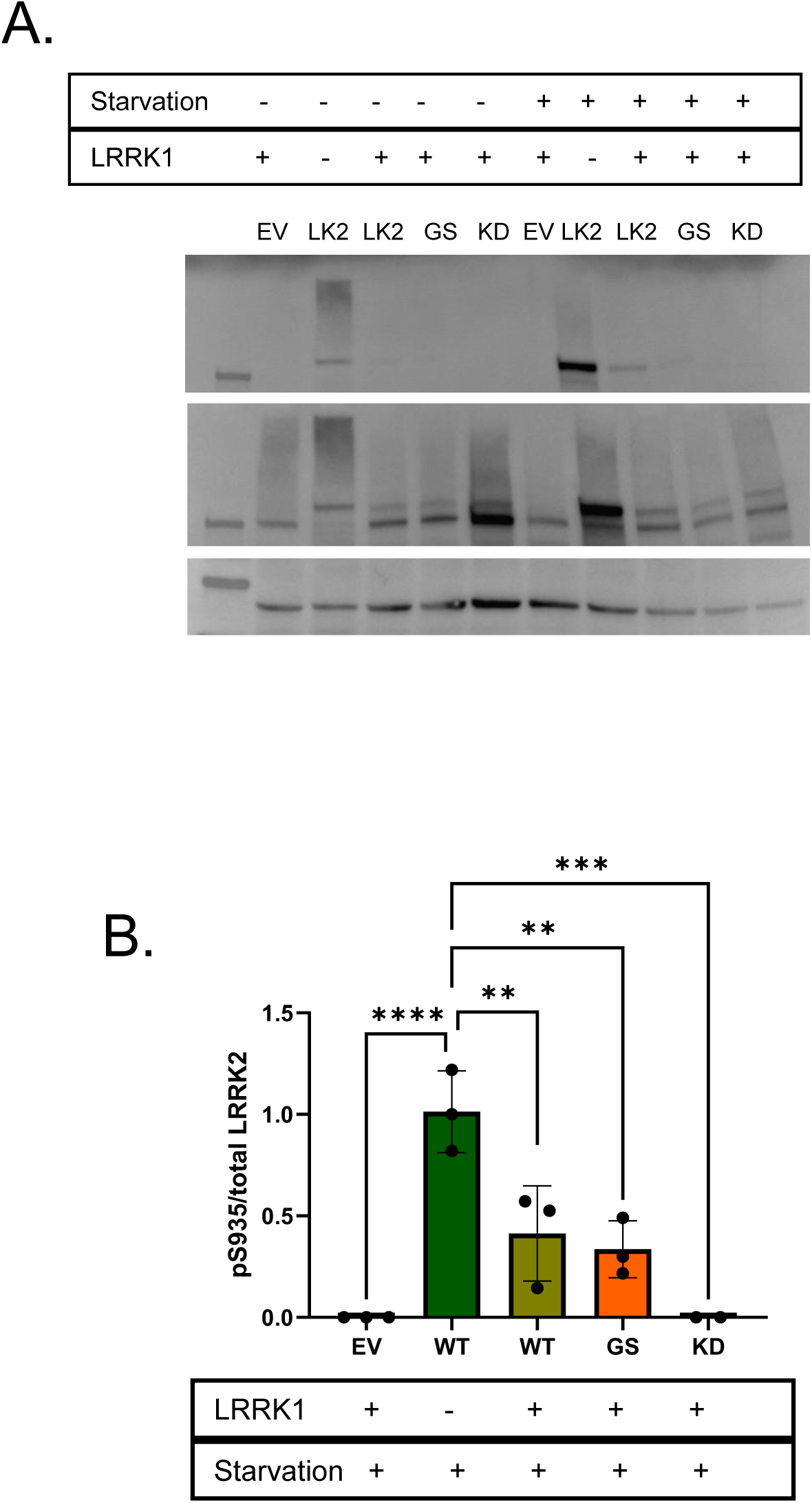
LRRK1 reduces the phosphorylation of LRRK2 at S935. (**A**) Western blot shows phosphorylation of LRRK2 at S935 under normal and starvation conditions. (**B**) Quantification of pS935 signal under starvation condition (n=3, 1-way ANOVA; Dunnett’s multiple comparisons test, *** p=0.0002).

### LRRK2 increases LRRK1 activity

Next, we investigated the effect of LRRK2 on LRRK1 kinase activity. Since Rab7 is a well-characterized substrate of LRRK1 but not LRRK2 (S1. C), we used it as a readout of LRRK1 activity. In cells expressing LRRK1, LRRK2, and Rab7, we observed a significant increase in Rab7 phosphorylation when LRRK2 was expressed. Notably, this enhancement occurred even in the presence of a kinase-dead LRRK2 mutant, indicating that the effect is independent of LRRK2 kinase activity (Fig 5. A and B). These results suggest that LRRK2 enhances LRRK1-mediated phosphorylation of Rab7 through a non-catalytic mechanism, possibly by enhancing the interaction between LRRK1 and Rab7 or stabilizing LRRK1 in an active conformation. Rab7 is a key regulator of late endosome and lysosome dynamics and localizes to late endosomes and promotes their maturation, fusion with autophagosomes, and lysosomal degradation processes essential for autophagy and cellular homeostasis. Given this role, we next studied how the LRRK1/LRRK2 interaction influences lysosomal function.

**Fig 5.**
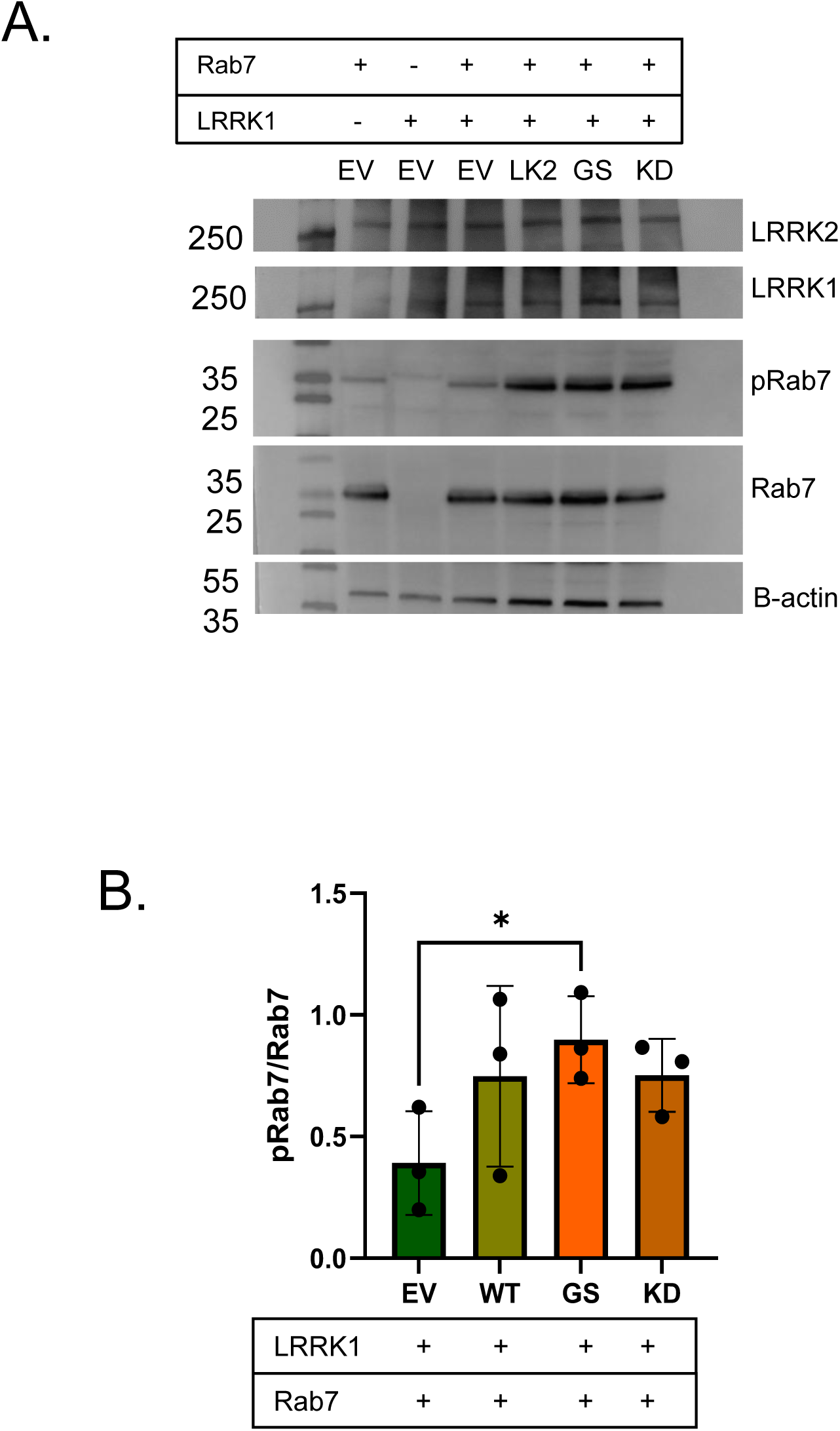
LRRK2 increases the activity of LRRK1 under normal conditions. (**A**) Western blot shows phosphorylation of Rab7 under normal and starvation conditions. (**B**) Quantification of A (n=3, Two- tailed t-test, P= 0.0343).

### LRRK2/LRRK1 interaction decreases lysosomal exocytosis

To examine the role of LRRK2 and LRRK1 in lysosomal size, we expressed Rab7-EGFP in cells transfected with WT LRRK2, G2019S, or KD variants. After 2 hours of incubation with Lysotracker, the Lysotracker signal was colocalized with Rab7-EGFP (S2. A), indicating that Rab7 is recruited to late endosomes and lysosomes.

Analysis of lysosomal size based on Rab7 signal showed that cells expressing LRRK2 G2019S exhibited significantly larger lysosomes compared to controls, whereas cells expressing LRRK2 KD showed smaller lysosome diameter (S2. A and B). On the other hand, cells expressing WT LRRK1 showed significantly smaller lysosomes (S2. C and D). Given that lysosomal size directly influences exocytosis, we next investigated the effect of these proteins on lysosomal exocytosis. We transfected cells with LRRK2 alone, LRRK1 alone, or both LRRK1 and LRRK2 (WT, G2019S, or KD). Lysosomal exocytosis was quantified by flow cytometry using an antibody against LAMP1, a lysosomal membrane protein that becomes exposed on the cell surface upon exocytosis. We found that cells expressing LRRK1 alone significantly increased lysosomal exocytosis. In contrast, cells expressing LRRK2 alone had no detectable effect. However, co-expression of LRRK1 with either WT LRRK2 or G2019S led to a reduction in exocytosis compared to LRRK1 alone. Interestingly, cells co-expressing LRRK1 with LRRK2 KD displayed a significant increase in lysosomal exocytosis (Fig 6. A and B). These results indicate that the LRRK2/LRRK1 interaction negatively regulates lysosomal exocytosis, and this inhibition is dependent on LRRK2 kinase activity.

**Fig 6.**
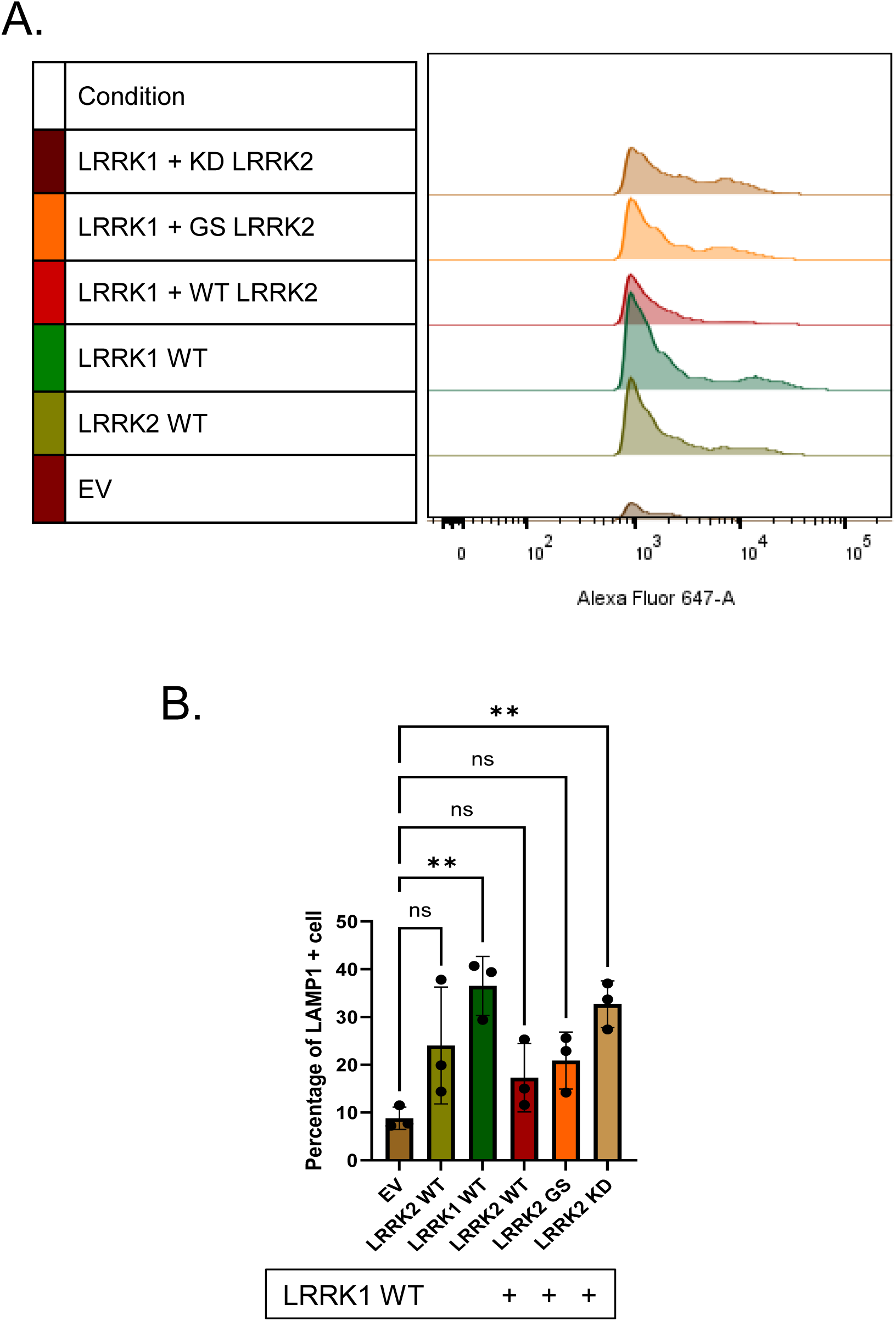
LRRK2/LRRK1 interaction decreases lysosome exocytosis. (**A**) Flow cytometry histograms for LAMP1 on the surface of HEK293T cells expressing EV, WTLRRK1, or WTLRRK1 with different variants of LRRK2. (**B**) Data represent the percentage of cell with LAMP1 signal (n=3, 1-way ANOVA; Dunnett’s multiple comparisons test, ** p=0.0049).

## Discussion

Mutations in *LRRK2* are associated with both familial and sporadic forms of PD. Although LRRK1 shares significant structural homology with LRRK2, it has limited established links to PD (32–34). Early structural studies suggested a potential interaction between LRRK1 and LRRK2 (28,33). However, the absence of strong evidence linking LRRK1 directly to PD has limited interest in studying this protein in the field of PD research. In this study, we used PLA to show the interaction between LRRK1 and LRRK2. This interaction was further validated using BiFC experiments. Given their structural similarity, yet distinct functional roles particularly in activation mechanisms and cellular regulation, we investigated how these proteins respond to changes in cellular nutritional status, including optimal nutrition and starvation, conditions known to trigger autophagy. We found that cellular nutrient conditions significantly influence the phosphorylation status of LRRK2, which is an indirect reflection of the LRRK2 activity. The activity of LRRK2 is regulated by Rab12 (35), which controls intracellular amino acid levels by modulating the trafficking of the plasma membrane amino acid transporter PAT4 from the cell surface to lysosomes (36). Phosphorylation at S935 primarily reflects a conformational change in LRRK2, which is unfavourable for LRRK1/LRRK2 interaction. Interestingly, our results indicate that the interaction between LRRK1 and LRRK2 is not dependent on LRRK2 kinase activity. However, the reduction in their interaction under starvation conditions is kinase-dependent. The kinase-dead mutant of LRRK2 shows minimal reduction in interaction upon starvation, whereas the pathogenic G2019S mutant exhibits the most pronounced decrease in signal. The LRRK2 kinase domain is a dynamic allosteric region, and the G2019S mutation lies within its activation loop. This mutation increases LRRK2 kinase activity by 2–3 fold, likely by stabilizing the active conformation of the kinase (37–40). This altered conformation may not only affect the enzymatic activity but also influence the protein-protein interaction, and that explains the reduction of LRRK1 and LRRK2 interaction under nutritional stress conditions. Our data reveal that the interaction between LRRK2 and LRRK1 induces a conformational change in LRRK2, evidenced by a significant decrease in phosphorylation at Ser935, favouring the kinase-inactive state. Conversely, LRRK1 kinase activity is enhanced under conditions of LRRK1/LRRK2 interaction, as indicated by increased phosphorylation of Rab7, a well-established substrate of LRRK1. However, it remains unclear whether this increase in Rab7 phosphorylation results from an enhancement in LRRK1 intrinsic enzymatic activity or from altered substrate accessibility due to changes in the LRRK1/Rab7 interaction kinetics.

Phosphorylated Rab proteins are retained at membrane compartments, including the lysosomal surface (41). When LRRK2 is recruited to membranes, phosphorylated Rab substrates accumulate at these sites (41–43). This accumulation can be counterbalanced by the phosphatase PPM1H, which dephosphorylates Rab proteins and facilitates their release from membranes, thereby restoring normal trafficking dynamics (44). In the context of LRRK2/LRRK1 interaction, hyperphosphorylated Rab7 accumulates on the lysosomal membrane, leading to a depletion of the inactive (non-phosphorylated) Rab7 pool. This imbalance disrupts normal lysosomal function and impairs processes dependent on dynamic Rab7, including endolysosomal trafficking, lysosome motility, and fusion events. Given that Rab7 is a master regulator of late endocytic trafficking, transport from late endosomes to lysosomes, lysosomal motility, clustering, and fusion at the perinuclear region (45,46). Hence, the disturbance of the Rab7 cycle leads to deregulation of cellular homeostasis.

Lysosomal dysfunction is a well-established hallmark of PD (7). Although the direct link between LRRK1 and PD remains unclear, its role in lysosomal regulation is well recognized. LRRK1 regulates lysosomal distribution, acidification, and protease exocytosis in osteoclasts, thereby controlling bone resorption (47). Our study clearly demonstrates that LRRK1 expression leads to a significant reduction in lysosomal size in contrast to cells expressing WT LRRK2 or the pathogenic G2019S mutant, which both exhibit enlarged lysosomes.

Lysosomal size, number, and positioning are dynamically regulated by a network of factors, including ER–lysosome contact sites, actin cytoskeleton, tethering and coat proteins, calcium signaling, phosphoinositides, V-ATPase activity, and mTORC1 signaling, which all influence lysosomal fission and fusion (48). Importantly, lysosomal size inversely correlates with lysosomal exocytosis: smaller lysosomes are more prone to exocytosis, a process critical for the extracellular release of undegraded material, including aSyn, particularly under conditions of lysosomal stress (e.g., after treatment with lysosomal-damaging agents or V-ATPase inhibitors). This mechanism has been proposed as a neuroprotective pathway in neurons (49).

Consistently, our data show that LRRK1 enhances lysosomal exocytosis, a phenotype that aligns with LRRK1 known role in bone remodelling. In contrast, LRRK2 expression does not promote exocytosis. Interestingly, LRRK2 expression, whether WT or G2019S, suppresses the exocytic activity induced by LRRK1, suggesting a functional antagonism between the two proteins. Our findings highlight the importance of studying the cellular mechanisms of LRRK1 in understanding PD. LRRK1 has been shown to play a fundamental role in regulating osteoclasts and bone mass through regulating lysosomal function, a key process directly linked to PD. A deeper understanding of this pathway could reveal a novel therapeutic target for developing DMTs.

## Methods

### Plasmids

The production of WT human LRRK2, G2019S, VN-LRRK2 and VC-LRRK2 was previously described (50). The kinase dead LRRK2 mutation (D1994A) was introduced by site-directed mutagenesis (Agilent Technology, CA, USA). The WT LRRK1 cDNA was a gift from Prof. Stefan Knapp, Goethe University Frankfurt.

For generating the LRRK1-VN fusion, the LRRK1 WT sequence was amplified by PCR (using the primers 5’GGGGCTTATGGCTGGCATGTCGCAAAGACCCCCCAGCATGTACTGGT GTGTGGGG3’ and 5’GGGGCTTAAGCTTATGGCTGGCATGTCG C AAAGACCCCCC AGCATGTACTGGTGTGTGGGG3’), then digested using the restriction enzymes AflII/XbaI, and subcloned into the same sites in the vector containing the VN cDNA. Positive clones were confirmed by restriction digestion and sequencing. For generating the LRRK1-VC construct, a similar approach was followed using the following primers: Forward 5’GGGGGGCTAGCATGGCTGGCATGTCGCAAAGACCCCCCAGCATGTACTG GTGTGTGGGGCCGG3’ and Reverse 5’GGGGGTCGACCCTTCTCTTGCGAGT GCAAGCCTCCAGCCGGCCCAGCTCC3’. PCR products were digested with NheI/XbaI enzymes, and subcloned into the same sites in the vector containing the VC cDNA.

### Cell cultures

The human embryonic kidney (HEK293) cells were grown in Dulbecco’s Modified Eagle’s Medium (DMEM) containing 4.5 g/L glucose (PAN Biotech), supplemented with 10 % (v/v) fetal calf serum (FCS) and 1 % (v/v) penicillin/streptomycin, and 1% non-essential amino acids (NEAA). The cells were maintained at 37 °C in a humidified atmosphere containing 5 % CO₂ under standard cell culture conditions.

### Immunocytochemistry (ICC)

Cells were fixed with 4% paraformaldehyde in DPBS for 10 min and permeabilized with 0.1% Triton X-100 in DPBS for 10 min. Cells were blocked in 5% BSA and incubated with primary antibodies (rabbit anti-FLAG, F7425 from Sigma; mouse anti-V5, R960-25 from Invitrogen) overnight at 4 °C. After three washes (10 min each) with DPBS, cells were incubated with the appropriate secondary antibodies for 1 h at room temperature. Cells were stained with DAPI for 5 min, and coverslips were mounted onto glass slides using Fluoromount-G (Life Technologies–Invitrogen). Slides were allowed to dry and stored at room temperature until imaging and analysis.

### Western blotting

Cells were lysed in RIPA buffer (50 mM Tris-HCl, pH 8.0; 150 mM NaCl; 0.1% SDS; 1% NP-40; 0.5% sodium deoxycholate; 2 mM EDTA) supplemented with protease and phosphatase inhibitors for 30 min on ice. Protein concentration was determined, and 30 µg of total protein was denatured in 5× loading buffer and separated on a 4–20% precast polyacrylamide gel (Bio-Rad). Proteins were transferred to nitrocellulose membranes using a Trans-Blot Turbo system (Bio-Rad). Membranes were blocked in 5% BSA in TBS-T for 1 h at room temperature and incubated with primary antibodies (rabbit LRRK2 (ab133474), rabbit LRRK2 pS935 (ab133950) and rabbit Rab7 pS72 (ab302494) from abcam, rabbit Rab7 (sc-10767) from Santa Cruz, rabbit flag (F-7425), and mouse beta-actin (A5441) from Sigma) overnight at 4 °C. After washing, membranes were incubated with HRP-conjugated secondary antibodies (anti-mouse or anti-rabbit from Amersham) for 1 h at room temperature. Signal was detected using enhanced chemiluminescence (ECL; Millipore), imaged using a Fusion system (Vilber Lourmat), and quantified using Fiji software.

### Proximity Ligation Assay (PLA)

PLA was performed according to the manufacturer’s instructions (Duolink®). Cells were fixed 48 h post-transfection with 4% paraformaldehyde, washed with DPBS, and permeabilized with 0.1% Triton X-100 for 10 min. After washing with 1× Wash Buffer A, cells were blocked in Duolink® Blocking Solution for 1 h at 37 °C. Primary antibodies (rabbit anti-FLAG, F7425; mouse anti-V5, R960-25) were diluted 1:1000 in Duolink® Antibody Diluent and incubated overnight at 4 °C. Cells were washed and incubated with PLA probes (1:5 dilution) for 1 h at 37 °C. Ligation was performed using ligase (1:40) for 30 min at 37 °C, followed by amplification using polymerase (1:80) for 100 min at 37 °C in the dark. After final washes (Wash Buffer B and 0.01× Wash Buffer B), coverslips were mounted using Duolink® In Situ Mounting Medium with DAPI and stored at 4 °C. Imaging was performed using a Zeiss Observer Z1 microscope (Carl Zeiss).

### Bimolecular Fluorescence Complementation (BiFC)

Cells were transfected 24 h after plating using Metafectene (Biontex) with plasmid DNA (2 µg per well; ratio 1:1). At 24 h post-transfection, medium was replaced with either Earle’s Balanced Salt Solution (EBSS; Gibco) or complete medium to induce starvation or maintain control conditions, respectively. Cells were incubated for 24 h before analysis.

For flow cytometry, cells were trypsinized, neutralized with complete medium, and pelleted by centrifugation. Cell pellets were resuspended in DPBS containing 2% FBS and 0.1% propidium iodide. Data were acquired using a Guava EasyCyte flow cytometer (Millipore), recording 5,000 events per sample.

### Lysosomal exocytosis

Cells were harvested 48 h post-transfection in cell sorting solution (2% FCS in PBS) and blocked in 1% BSA for 20 min. Cells were incubated with primary antibody against Lamp1 (sc-20011; 1:200 dilution in 1% BSA) for 20 min on ice, washed twice with cold 2% FCS in PBS, and incubated with Alexa Fluor 633–conjugated goat anti-mouse IgG secondary antibody (1:500; Life Technologies–Invitrogen) for 20 min on ice.

Cells were washed, resuspended in 300 µL cold FACS buffer, and analyzed by flow cytometry. A total of 10,000 events were recorded per sample using a BD LSR Fortessa X-20.

## Statistical analysis

Data were obtained from at least three independent experiments and are shown as mean values ± standard deviation (SD). Two-group comparisons were performed using Student’s *t* test, multiple-group comparisons were performed using ANOVA with Šídák’s or Dunnett’s multiple comparisons test. *p*□<□0.05 was considered statistically significant (\**p* < 0.05; \*\**p* < 0.001; \*\*\**p* < 0.0001). Statistical analyses were performed in GraphPad Prism.

## Acknowledgments

We thank the Core Facility for Cell Sorting of the University Medical Center Göttingen (Germany). M.M. is supported by DAAD Research Grants - Doctoral Programmes in Germany, 57552340. T.F.O. is supported by the Deutsche Forschungsgemeinschaft (DFG, German Research Foundation) under Germany’s Excellence Strategy, EXC 2067/1- 390729940, and by SFB1286 (B8).

## Author contributions

T.F.O. and M.M. designed the study. E.G. and M.M. prepared the plasmids. C.F. provided technical support. M.M. and M.H. performed the experiments. C-A.B. provided scientific advice. M.M. and T.F.O. wrote the manuscript. All authors revised and approved the manuscript.

## Ethics Statement

This study did not require ethical approval.

## Data Availability Statement

All data are available from the corresponding author upon reasonable request.

---

**Supplementary Fig 1.**
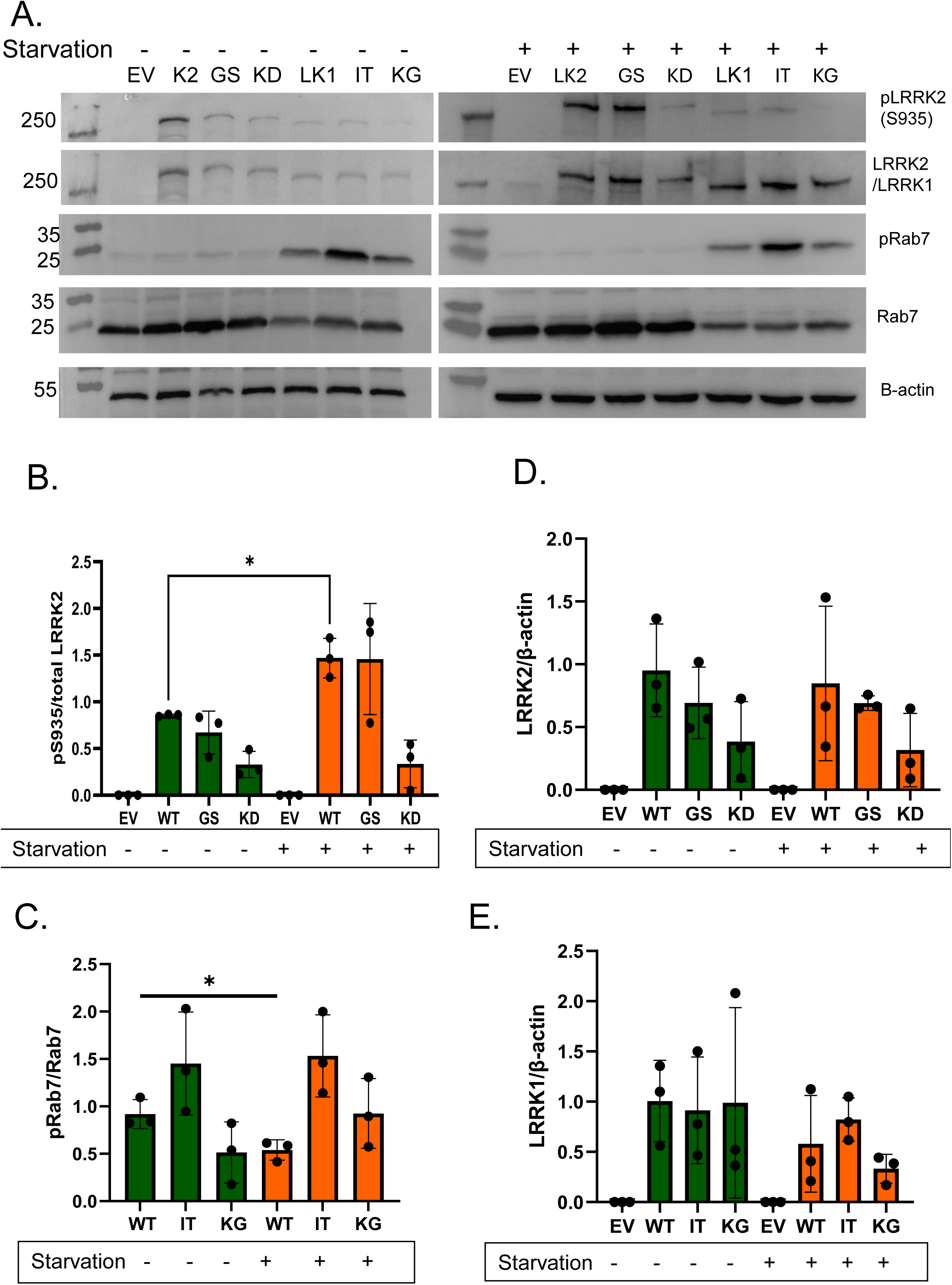
The phosphorylation of Rab 7 decreases under starvation conditions. (**A**) Western blot shows phosphorylation of Rab7 and LRRK2 at S935 under normal (Left) and starvation conditions (Right). (**B-E**) Quantification of A (n=3, T-test, P= 0.0370 for pS935 and P= 0,0296 for pRab7).

**Supplementary Fig 2.**
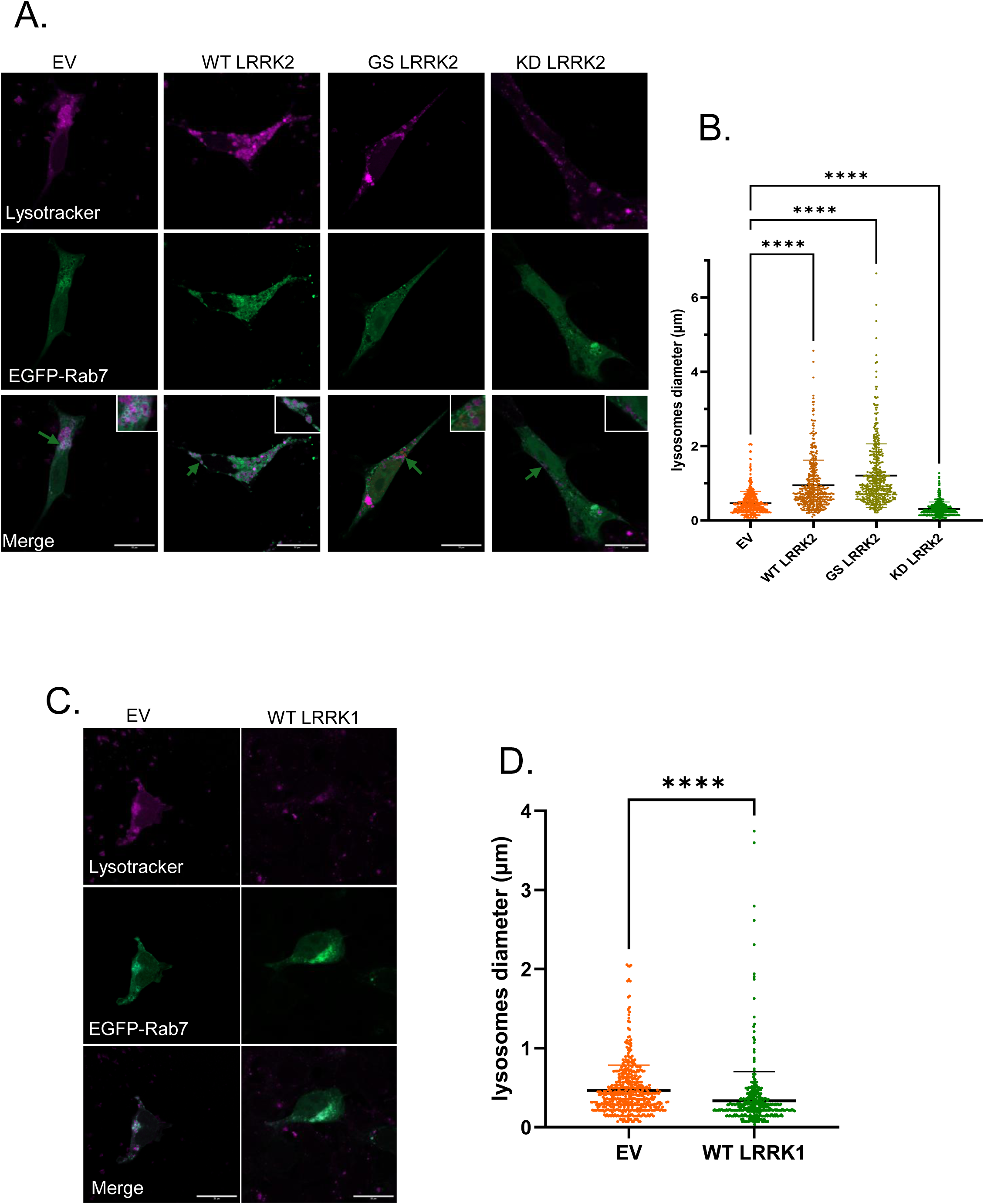
Increased kinase activity in LRRK2-expressing cells associates with larger lysosome diameters. (**A**) Confocal images show the lysosome signals in cells expressing EV and different variants of LRRK2. (**B**) Data represent the lysosome diameter in A (n=3, 1-way ANOVA; Dunnett’s multiple comparisons test, **** p<0.0001; at least 20 lysosomes for 10 cells per group across one independent experiment were analysed). (**C**) Confocal image shows the lysosome signals in cells expressing EV and WT LRRK1. (**D**) Data represent the lysosome diameter in cells expressing EV and WT LRRK1. (Two- tailed t-test, **** p<0.0001, at least 10 cells per group across three independent experiments were analysed).

